# Properties of cell signaling pathways and gene expression systems operating far from steady-state

**DOI:** 10.1101/334649

**Authors:** Juan Pablo Di Bella, Alejandro Colman-Lerner, Alejandra C. Ventura

## Abstract

Ligand-receptor systems, covalent modification cycles, and transcriptional networks are basic units of signaling systems and their steady-state properties are well understood. However, the behavior of such systems before steady-state is poorly characterized. Here, we analyzed the properties of the input-output curves for each of these systems as they approach steady-state. In ligand-receptor systems, the EC_50_ (concentration of the ligand that occupies 50% of the receptors) is higher before the system reaches steady-state. Based on this behavior, we have previously defined PRESS (for pre-equilibrium sensing and signaling), a general “systems level” mechanism cells may use to overcome input saturation. Originally, we showed that, given a step stimulation, PRESS operates when the kinetics of ligand-receptor binding are slower than the downstream signaling steps. Now, we show that, provided the input increases slowly, it is not essential for the ligand binding reaction itself to be slow. In addition, we demonstrate that covalent modification cycles and gene expression systems may also operate in PRESS mode. Thus, nearly all biochemical processes may operate in PRESS mode, suggesting that this mechanism may be ubiquitous in cell signaling systems.

## Introduction

Cells detect input signaling molecules using receptors, proteins usually located on the cell surface embedded in the plasma membrane. Activated receptors then transmit the signal to the interior of the cell through a series of downstream processes that typically lead to changes in gene expression, resulting in an appropriate output response to the input. In a way, the system’s overall input-output curve summarizes its biological characteristics and function [1]. Several of the input-output characteristics are well appreciated [2]. For example, some input-output curves are graded, changing outputs moderately over a wide range of inputs; other are ultrasensitive, changing very rapidly from low to high output across a narrow range of inputs. The response of biological systems to changing environmental conditions is a dynamic process. The response time (the time to reach 50% of its value at steady state) of a given system depends on the input strength and duration, as well as on the structure and the kinetic parameters of the system under consideration [3]. These two notions, the shape of an input-output curve and the dynamics involved in reaching its steady-state, have not been properly integrated in the literature. The focus of the current work is on studying how an input-output curve evolves over time and how this evolution confers dynamic tunability to the signaling system.

In a ligand-receptor system, the EC_50_ (the concentration of the ligand that occupies 50% of the receptors) changes over time [4]. In the case of a single binding site, the EC_50_ is elevated at at early times, and it drops to its steady-state value as binding approaches equilibrium. At that time, the EC_50_ is equal to the dissociation constant of the ligand:receptor reaction. Thus, in effect, the dose-response binding curve *shifts* over time from right to left [4-6]. Note that a *shift* in the binding curve over time implies that a ligand-receptor system is sensitive (ie, a change in the input concentration elicits a change in the output) in different regions of ligand concentration at different times before steady-state [4]. Based on this property, we observed that when the ligand:receptor complex activates a downstream signaling component with a time-scale compatible with acting before the ligand:receptor binding process achieves equilibrium, this signaling component will use the information contained in the pre-steady-state binding curve, and thus produce an output with an EC_50_ shifted to the high ligand concentration region. Therefore, such a system provides distinguishable responses to ligand concentrations that saturate the receptors at steady-state. We termed this “systems level” mechanism PRESS, for pre-equilibrium sensing and signaling, and we showed its potential role in yeast directional polarization in response to pheromone gradients [4].

We have coined the term *shift* for the property of systems that change their EC_50_ over time and refer to systems that exhibit this property *shifters*. A *shift* mathematically means that the EC_50_ has to be a time dependent function. In a ligand-receptor system, this *shift* is possible because the time to reach binding equilibrium depends inversely on the ligand concentration: the higher the ligand concentration, the faster the equilibrium is reached, see Figure 1B and D. Thus, at early times after addition of the ligand, the binding process will be further from equilibrium at the lowest ligand concentrations, see Figure 1C. This will result in a higher EC_50_ value than at equilibrium and a steeper binding curve (i.e., less graded, Figure 1C and Figure 1E). Curve steepness is often quantified using the Hill coefficient, n_H_, defined as the log(81)/log(EC_90_/EC_10_), where EC_90_ and EC_10_ are the inputs required to elicit 90% or 10% of the maximum output.

**Figure 1:**
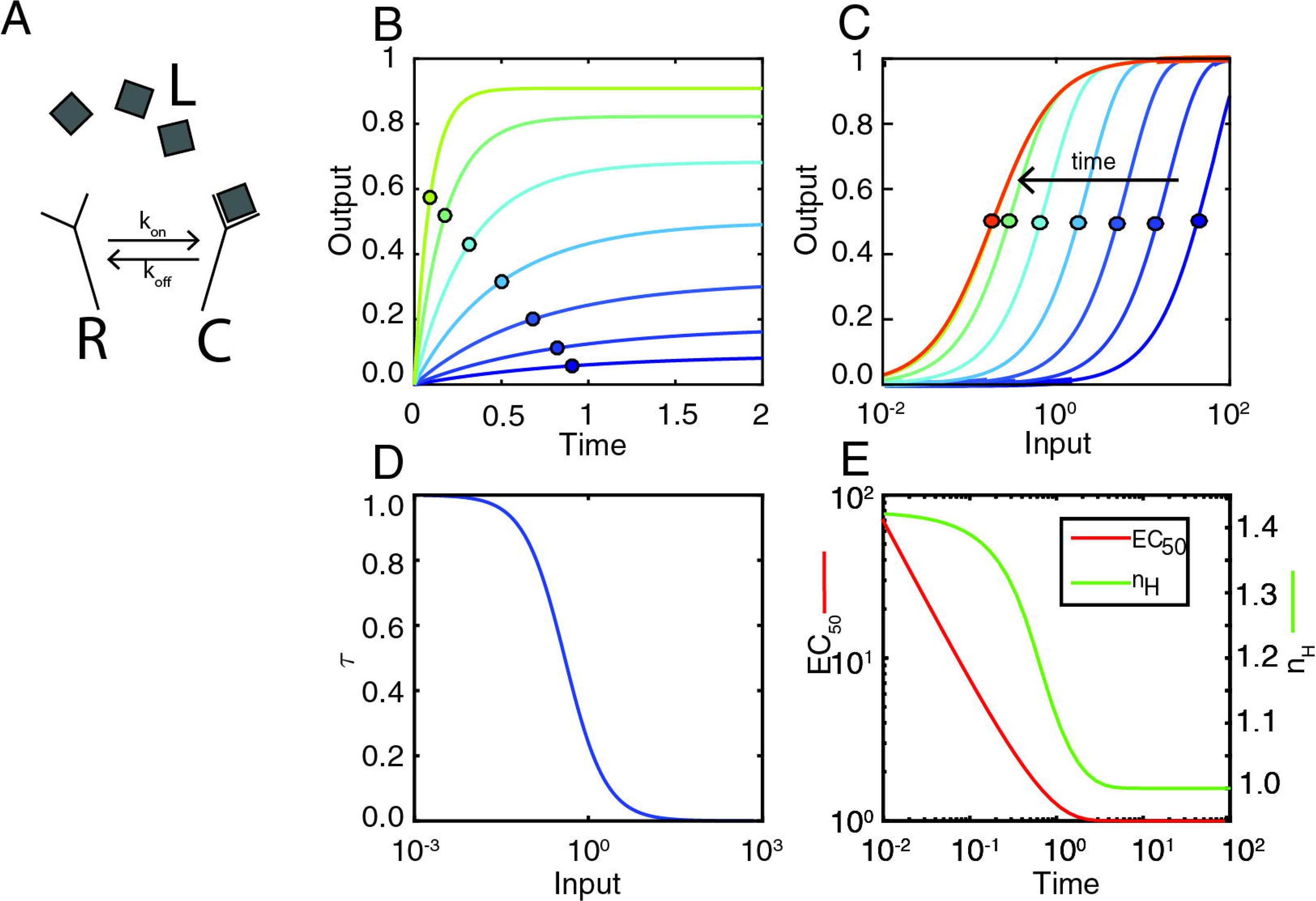
Characterization of a prototypical shifter: a ligand-receptor reaction. **A** Ligand-Receptor scheme, ligand binds free receptor reversibly with rate constants k_on_ and k_off_. **B**. Output (C, ligand-receptor complex) vs. time, for different values of input (color-coded using a heat-map scale). Circles over the curves mark output at time τ (time at which output is 63.2% of its steady-state value). **C**. Input-output curves obtained at different times (color-coded using a heat-map scale). Circles mark the EC_50_. **D**. τ vs. input. **E**. EC_50_ (red) and nH (green) vs. time.

Here, we asked if the concept of PRESS may be applied to processes that do not reach thermodynamic equilibrium, such as ligand receptor binding, but do reach a steady state, such as covalent modification (CM) cycles and gene expression systems. We found that both can indeed be shifters. Furthermore, we show that by introducing dynamics to the input it is possible to extend PRESS to fast signaling components, effectively extending this property to all signaling motifs and most parameter sets.

## Results

### Ligand-receptor systems revisited: a slowly increasing input controls the *shift*

Under physiological conditions, inputs (e.g., concentrations of growth factors or nutrients) are likely to change gradually rather than in a stepwise manner [7]. Therefore, to determine how PRESS is altered when the input has dynamics, we considered the case of a ligand *L* that increases exponentially and interacts with a receptor *R* to form an activated complex *C*, with binding and unbinding rates *k*_*on*_ and *k*_*off*_, respectively. We assumed that total amount of receptor is constant *R*_*T*_ = *R* + *C*, and that the concentration of free *L* is not depleted significantly by the binding reaction, so that *L∼L*_*T*_ (total amount of ligand). Thus, we described the system by the following equations:

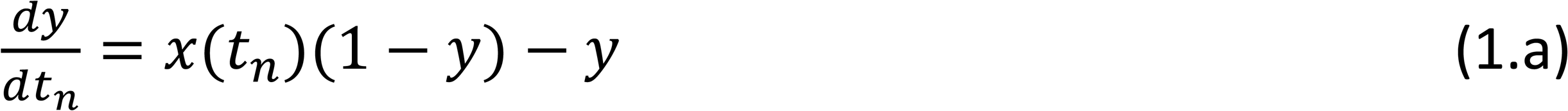

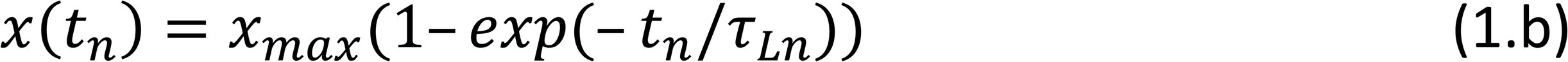

where *x*(*t*_*n*_) = *L*(*t*_*n*_)*/K*_*d*_ is the amount of ligand relative to the dissociation constant for the binding-unbinding reaction (*K*_*d*_ = *k*_*off*_*/k*_*on*_), *y* = *C/R*_*T*_ is the amount of ligand-receptor complex relative to the total amount of receptors, and *t*_*n*_ = *t/t*_*ref*_ is the time expressed in units of a reference time *t*_*ref*_ *=* 1*/k*_*off*_. We further assumed that the amount of ligand increases following an exponential function characterized by a maximum value *L*_*max*_ and a characteristic time *τ*_*L*_, resulting in *x*_*max*_ = *L*_*max*_*/K*_*d*_ and *τ*_*Ln*_ = *τ*_*L*_*/t*_*ref*_, which connects the time-scale associated with the ligand accumulation (*τ*_*L*_) with that of the binding-unbinding process (*t*_*ref*_ *=* 1*/k*_*off*_).

We solved equation (1a-b) using three input dynamics, much faster (nearly a step increase), of the same order or much slower than the binding reaction timescale (*τ*_*L,n*_ *=* 0.01, 1 and 100, respectively), for a range of input *x*_*max*_ (Figure 2 A,B,C). These simulations showed two important results. First, the response *y* reached steady-state faster when the input was larger (Figure 2 D,E,F), even though the time for the input *x* to reach *x*_*max*_ was modeled so that it was independent of its magnitude. Consequently, the *y* vs *x*_*max*_ plot shows a clear *shift* in the binding curve over time (Figure 2 G,H,I), with a lower bound of EC_50_ = 1. Second, the dynamics of the stimulus affected the speed of the *shift*: larger values of *τ*_*Ln*_ resulted in slower *shifts*. That is, a slow rising input caused a slow *shift* (Figure 2 J), providing a longer time to a pathway downstream to operate in PRESS mode.

**Figure 2:**
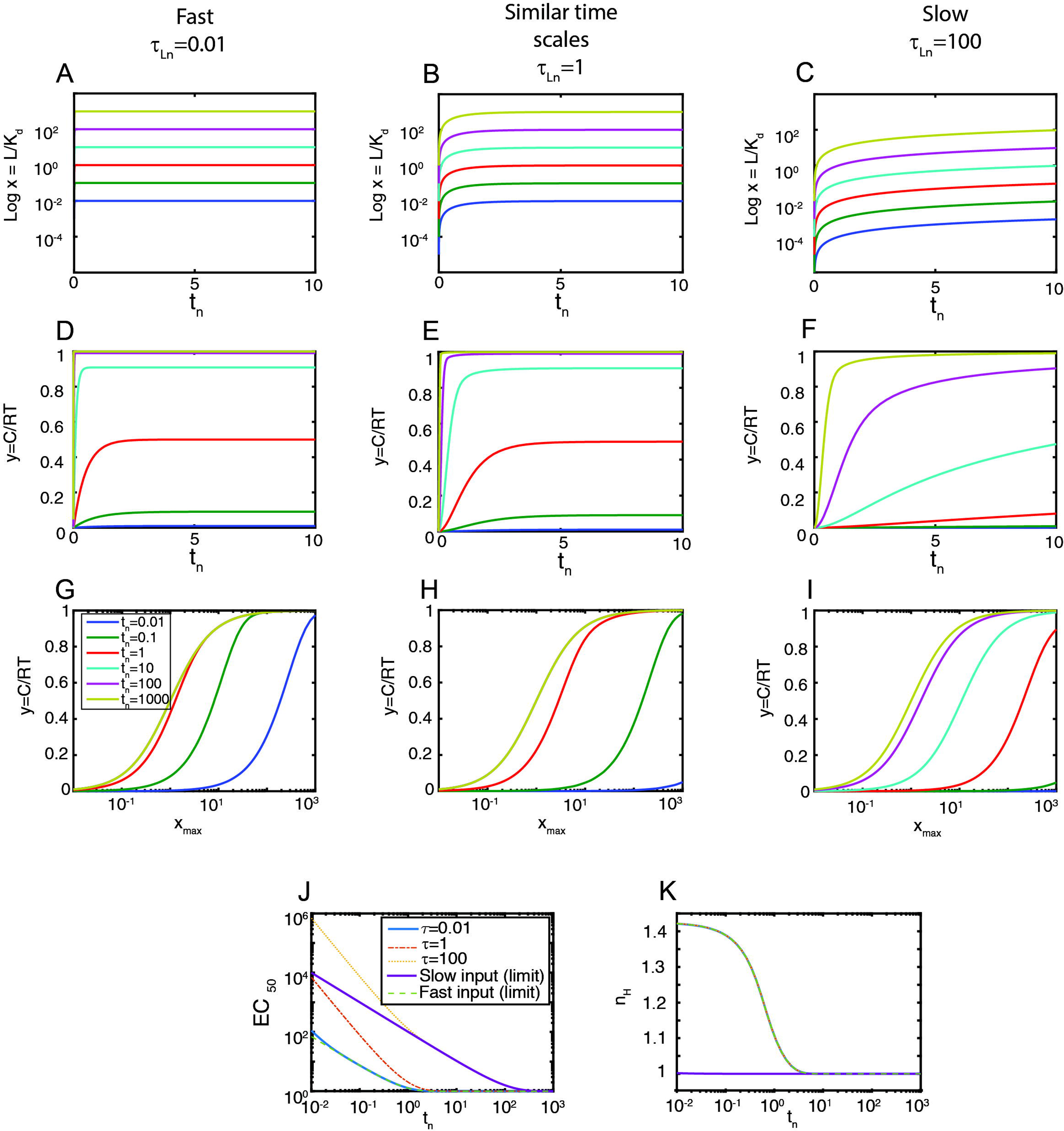
Ligand-receptor system. Left panels τ_Ln_ = 0.01 (fast stimulation), middle panels τ_Ln_ = 1, right panels τ_Ln_= 100 (slow stimulation). **A-C**. Stimulus vs. time (t_n_=t/t_ref_), colors indicate x_max_ = 0.01-0.1-1-10-100-1000. **D-F**. Response y vs. time (t_n_=t/t_ref_), colors indicate x_max_ as before. **G-I**. Input-output curves, response y vs. input (x_max_) for the indicated times. **J.** EC_50_ vs time for the indicated input dynamics. **K.** Hill coefficient (n_H_) vs time for the same conditions used in **J**. In **J** and **K**, the curves indicated as slow and fast input, respectively, were obtained with the corresponding analytical results in the SI. The fast input curve completely overlaps the τ = 0.01 one.

We also computed the Hill coefficient n_H_ for the three input dynamics. We found that the temporal profile for the three dynamics was the same (Figure 2K). This result indicates that the dynamics considered for the stimulus has no effect on the dynamics of n_H_.

To quantitatively understand the relationship between the timing of the *shift* and the time scales of the stimulus and receptor binding, we derived analytical results for two limit cases: fast increase in the input with slow binding (*τ*_*Ln*_ ≪ 1) and the converse, slow increase in the input with fast binding (*τ*_*Ln*_ ≫ 1) (see Supplementary Information). Particularly, in this last limit case, assuming that the ligand-receptor complex is in quasi-steady state with the value of the input *x* at each time, we obtained:

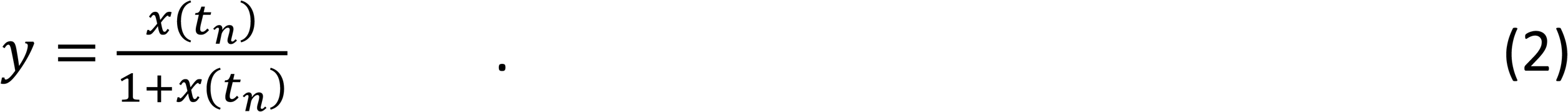

This equation is characterized by

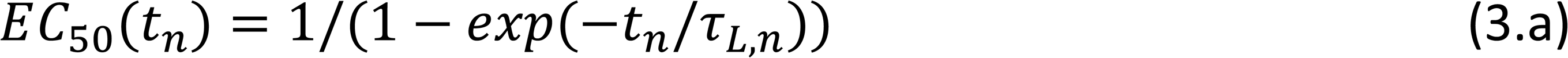

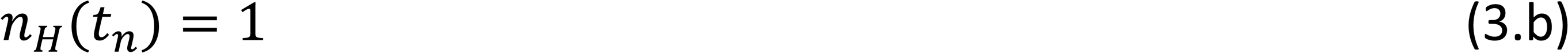

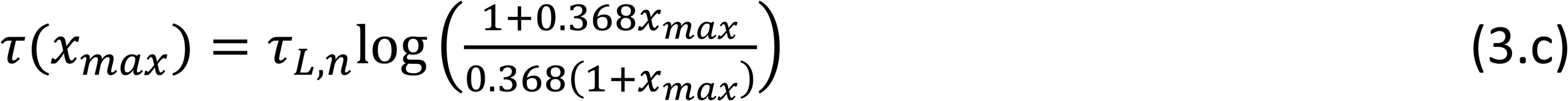

Here, equation (3.a) is a decreasing function of time, equation (3.b) shows a time independent n_H_, and (3.c) is a decreasing function of the input, (see Supplementary Information for details). These results indicate that in this limit case, the *shift* is conserved: for the convolved process *τ* is a decreasing function of the input and this causes the EC_50_ to be a decreasing function of time (Figure 2J). Moreover, the time-dependency of the EC_50_ is exactly one divided by the time-dependency of the input, an exponential function of time in the example considered. Thus, if the input increases slowly, the corresponding EC_50_ decreases slowly. Interestingly, and related to results in Figure 2K, we found that, for fast binding, n_H_ is constant over time, indicating that the dynamics of the Hill coefficient is governed by the binding reaction independently of the stimulus temporal profile.

In summary, in previous work we had noted that operation in PRESS mode required that the “sensing” ligand-binding reaction be slower than the downstream “signaling” component [4]. In that study, we considered a step-like stimulation, here corresponding to τ_Ln_ << 1. Our results here indicate (Figure 2) that the rising time of the input *τ*_*L,n*_ controls the speed of the shift of the input-output curves. Thus, by introducing dynamics to the rise in the concentration of the input, it is not essential any more that the sensing component be particularly slow. It is enough that the stimulus rises slowly relative to the downstream component, even in cases where the ligand-receptor interaction is fast.

### Covalent modification cycles

CM cycles consist of a substrate *S* that is activated into *S*^*^ by an enzyme through a post-translational modification such as phosphorylation and deactivated by another enzyme through the removal of that modification (Figure 3A). Here we studied their behavior pre-steady state in response to the accumulation of the activating enzyme *E*_*a*_, with constant deactivating enzyme *E*_*d*_. We used a mechanistic description as detailed in the SI. For this system, we considered that the input is x=(k_cat,a_ E_a_)/(k_cat,d_ E_d_), the ratio of the maximum velocities of the activating and deactivating enzymatic reactions, with catalytic rate constants k_cat,a_ and k_cat,d_; the output is y=S*/S_T_, the fraction of active substrate; and the two relevant parameters are K_a_= K_m,a_ / S_T_ and K_d_=K_m,d_/S_T_, the Michaelis-Menten constants (K_m_=(k_d_+k_cat_)/k_a_) relative to the total amount of substrate. In what follows, the time is expressed in units of a reference time t_ref_=S_T_/(k_cat,d_ E_d_), so that t_n_=t/t_ref_. (For a simplified Michaelis-Menten description, see SI).

**Figure 3:**
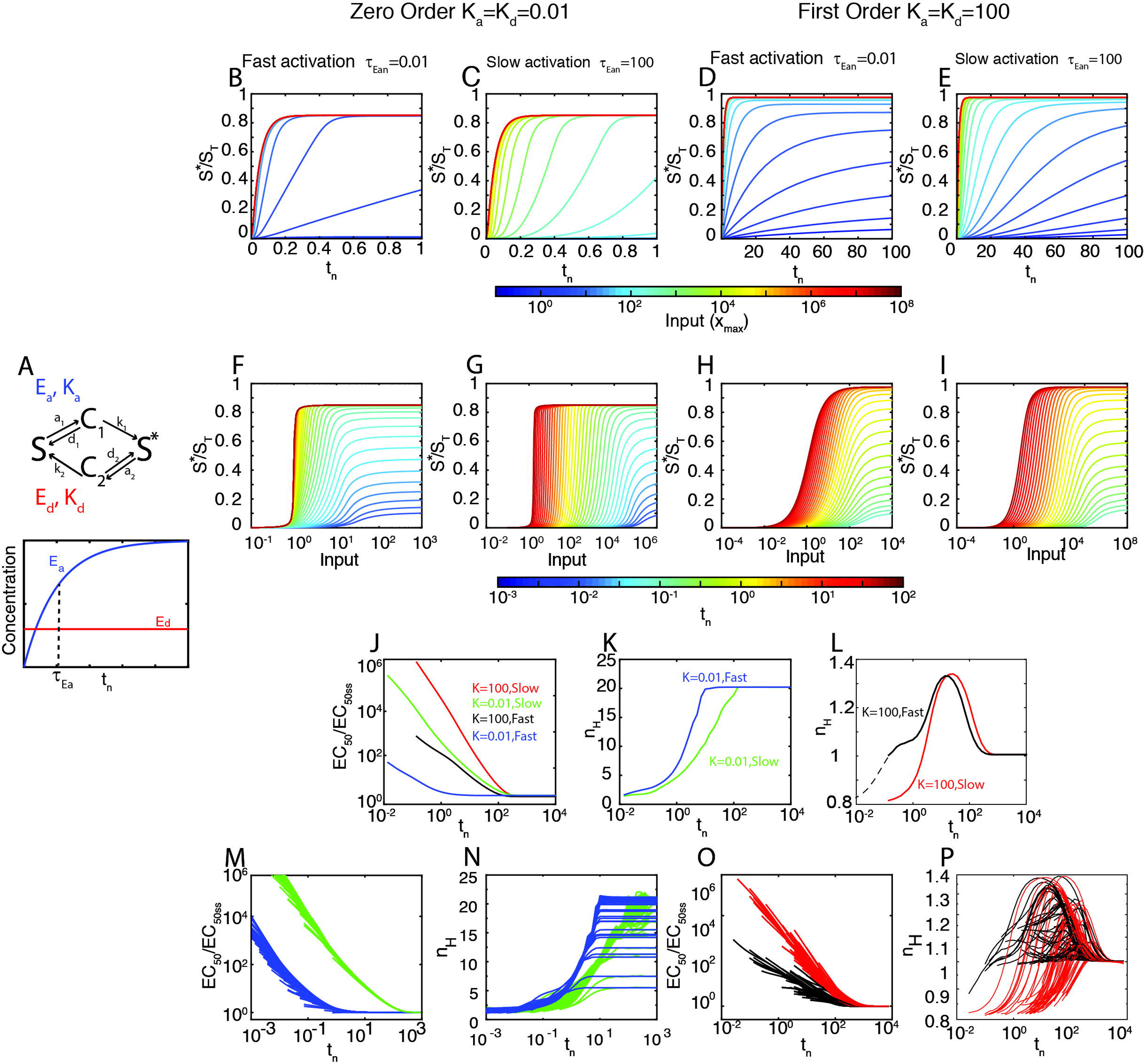
Covalent modification (CM) cycles are shifters. **A**. Top. Diagram of the enzymatic cycle. Bottom. Dynamics of E_a_ and E_d_ for exponential accumulation of E_a_. **B-E.** Time courses of y (=S*/S_T_) for different values of the input x_max_ (in a heatmap scale). **F-I.** Input-output curves at different times (in a heatmap scale). **B-I** correspond to 4 different examples, as indicated: K_a_=K_d_=0.01 (zero^th^ order regime, **B,C,F** and **G**) and K_a_=K_d_=100 (first order regime, **D, E, H** and **I**), each of them with τ_Ean_ = 0.01 (fast rising input, **B, D, F** and **H**) or τ_Ean_ = 100 (slow rising input, **C, E, G** and **I**). The complete list of parameters values for each example is included in the SI. **J-L.** EC_50_ (**J**) and n_H_ (**K** and **L**) vs time for the four examples considered in **B-I**. The inset in panel **L** contains earlier times for the case K=100, fast rising input). **M-P.** EC_50_ (**M** and **O**) and n_H_ (**N** and **P**) vs time for different parameter sets corresponding to the zero-order regime (**M** and **N,** in green and blue) or the first order regime (**O** and **P**, in red and black). For each scenario (K=100 slow, K=100 fast, K=0.01 slow, K=0.01 fast), 100 random parameter sets were scanned, see SI for details of the parameter scanning.

We further assumed that the activating enzyme increases following an exponential function, for example due to its accumulation after its expression is induced. This function is characterized by a maximum value E_a,max_ and a characteristic time τ_Ea_, resulting in:

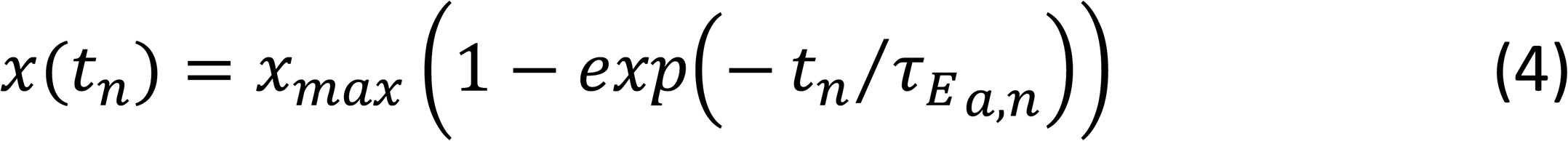

where 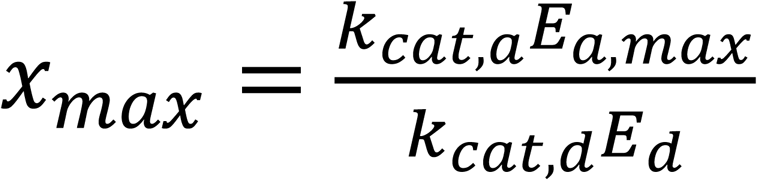 and 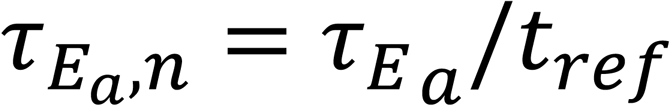.

Depending on the values of the Michaelis-Menten constants K_a_ and K_d_, two extreme regimes may be considered. In one regime, K_a_ and K_d_ are large (K_m,a_ >> S_T_ and K_m,d_ >> S_T_), and the kinetics of the reactions are first order (i.e., linear) with respect to the concentration of the substrate. In the other extreme regime, K_a_ and K_d_ are small (K_m,a_ << S_T_ and K_m,d_ << S_T_), the enzymes are saturated by their substrates, and therefore the kinetics of the reactions are zero-order with respect of the concentration of the substrates (only dependent on the amount of enzyme). In this second regime, the steady-state input-output curve is ultrasensitive [8], i.e., below a threshold input concentration, changes in input cause almost no change in output. However, past that threshold, a small change in input gives a much larger change in output. Systems with high ultrasensitivity thus behave as molecular switches.

We studied the system in these two extreme regimes (see methods for a detailed description of the parameter scanning), for the case in which the input x increases fast relative to the timescale of the covalent modification cycle (τ_Ea,n_ << 1), and thus it is essentially a step given by x(t_n_) = x_max_; and the case in which it is slow compared with the cycle (τ_Ea,n_ >> 1).

Figure 3 (B-L) shows simulations of example parameter sets of each of the four conditions (first-or zero-order, fast or slow input), as well as a parameter scan summarized in Figure 3 M-P (details in the Supplementary Information). Notably, the input-output curve *shifted* from right to left over time in all four conditions (Figure 3 F-I). Just as in case of the ligand-receptor system, when the stimulus increased slowly (Figure 3 C, E, G, and I), the leftward *shift* in the input-output curve shifted correspondingly slowly.

The above results thus show that CM cycles are shifters. However, we observed interesting differences between CM cycles working in zero-order and first order regimes. First, the *shift* was faster in zero-order, in the sense that the EC_50_ reached steady state values earlier (Figure 3J). This in turn indicates that there is less time for PRESS when the enzymes are saturated. Second, the n_H_ was biphasic in the first-order regime (Figure 3K) with an increase from about 0.85 to 1.35 and a later decrease towards 1. In stark contrast, in the zero-order regime n_H_ increased monotonically, from about 1 towards its final high steady-state value (∼5 to 22 in our simulations) (Figure 3L). This surprising result indicates that the well-known ultrasensitivity of this type of system is not immediately apparent but rather develops over time. In addition, that an ultrasensitive system is a shifter implies that the threshold input required to activate the switch moves over time. Thus, before reaching steady-state, much higher input concentrations would be needed to cross the threshold and flip the switch than at steady-state.

### Coupling shifters (I): a ligand-receptor reaction activating a covalent modification cycle

To study how the shifting property propagates in a system, we analyzed a toy system composed of the two shifter modules we have described so far: ligand-receptor coupled to a CM cycle (LR-CM) (Figure 4A). We implemented a full mechanistic model, including enzyme-substrate complexes. (See Supplementary Information for a complete description of the model). We defined two outputs: an upstream output *y =* (*RL* + *RLS*)*/R*_*T*_, the concentration of ligand-receptor in units of the total amount of receptor, and a downstream output *z* = *S*^*^*/S*_*T*_, the active substrate in units of the total amount of substrate. We then compared the times at which each output reached steady-state (*t*_*ssz*_ and *t*_*ssy*_) and the corresponding EC_50_ss, using different combinations of parameters so that the CM subsystem was either in the first or zero-order regime (see Methods). We made several observations. First, the LR-CM system as a whole was a shifter, since EC_50Z_ shifted from right to left over time (Figure 4C-D). Second, as expected for a linear system (ie, no feedback loops) [9], EC_50SSZ_ was always lower than EC_50ss_y (Figure 4 C), indicating that the range of EC_50_ spanned by the system was larger than that spanned by the first step alone. Third, in contrast to the curves of *y*, the amplitude of the curves of *z* increased over time. That is, at high input, the output was submaximal if the elapsed time was short enough. This is because at high input concentrations, ligand binding is faster than the steps downstream of receptor activation, and thus the speed of accumulation of z at those input concentrations is input independent. Fourth, *t*_*ssz*_ was always larger than *t*_*ssy*_ (Figure 4 C-E). Interestingly, we found that the ratio between the *t*_*ss*_ for the two outputs correlated well with the ratio of their *EC*_50_*ss* (Figure 4G): when the EC_50_ss are nearly identical (a particular case known as Dose-Response Alignment [9, 10], *t*_*ss*_ are also identical.

**Figure 4:**
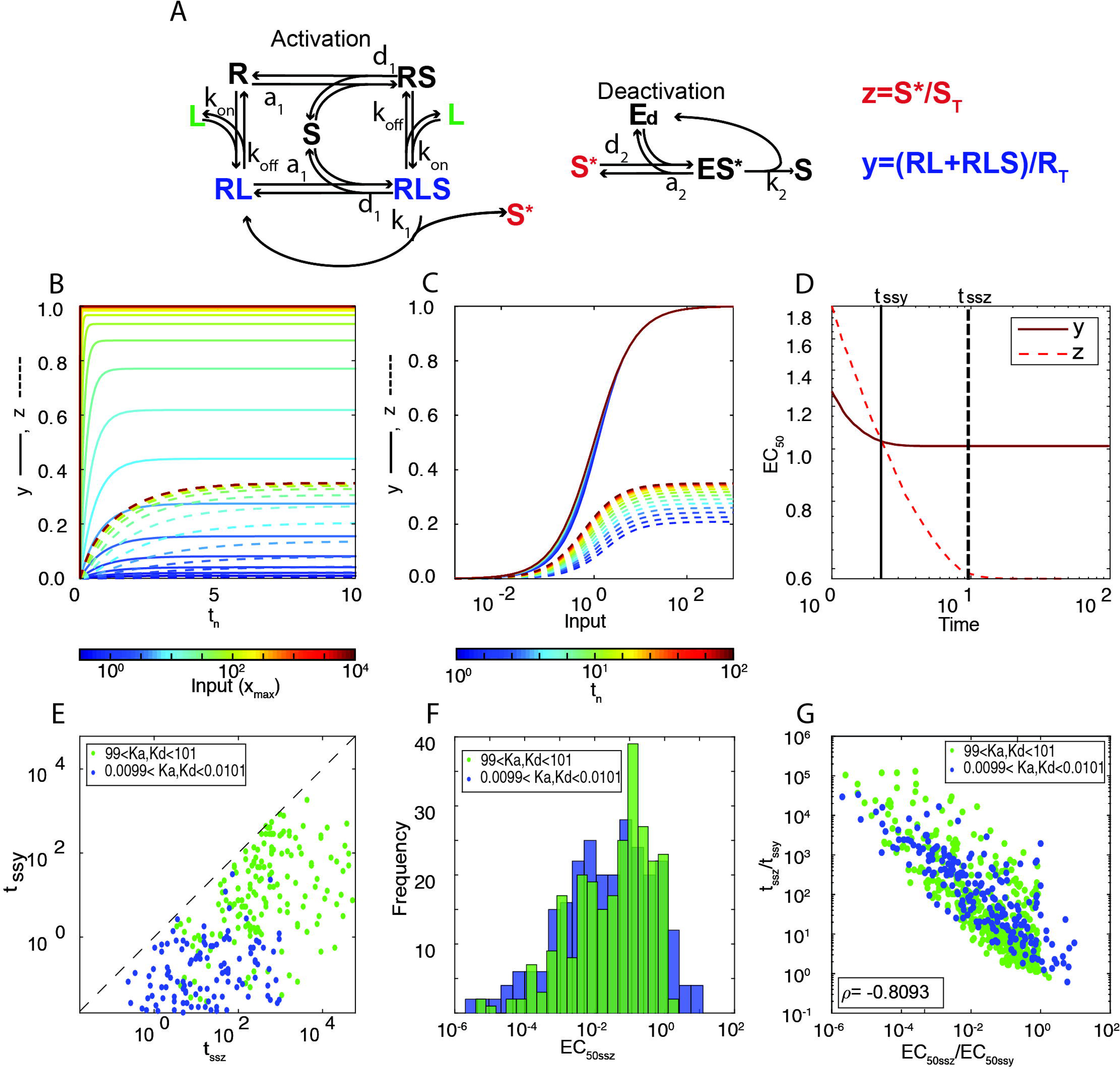
The LR-CM system is a better shifter than either in isolation. **A**. Reaction scheme of a biological network composed of a LR coupled to a CM cycle. The receptor has four possible states: unbound (R), bound to the ligand (RL), bound to the substrate (RS), and bound to both (RLS). RLS is also the enzyme-substrate complex that catalyzes S activation into S*. The enzyme E_d_ forms a complex with S*, ES* to deactivate it into S, completing the CM cycle. **B-C.** For an example parameter set, plot shows the time course and input-output curves (y=((RL+RLS)/R_T_), solid lines and z=(S*/S_T_), dashed lines) at different times distributed uniformly in log scale from t_0_ = 1 to t_f_ = 10^2^ (using a heatmap color scale). Each curve is normalized by its maximum value at steady-state. **D.** For the same example as in B and C, EC_50_ vs time for both outputs. Vertical lines mark the time t_ss_ at which each curve reached ss. Parameter set used in **B, C** and **D**: a_1_ = a_2_ = d_1_ = d_2_ = k_1_ = k_2_ = E_T_ = S_T_ = R_T_ = 1. Binding rates a_1_ and a_2_ are in units of ([time][concentration])^−1^, unbinding rates d_1_, d_2_, k_1_ and k_2_ are in units of [time]–1 and total amounts E_T_, S_T_ and R_T_ in units of [concentration]. **E**. t_ssy_ vs. t_ssz_ for 200 parameter sets sampled randomly within the ranges defined in Table S1 so that K_a_ [((d_1_+k_1_)/(a_1_*S_T_))] and K_d_ [((d_2_+k_2_)/(a_2_*S_T_))] fell in the ranges indicated. **F**. EC_50 SSZ_ for the 200 sets shown in E. **G.** The relative speed of the two modules t_ssz_/t_ssy_ negatively correlates with the relative sensitivity (EC_50ssz_/EC_50ssy_).

### Coupling shifters (II): a cascade of covalent modification cycles

In order to evaluate the shifting property in a concrete case, we considered the chain of events involved in oocyte maturation [11]. *Xenopus* oocytes convert a continuously variable stimulus, the concentration of the maturation-inducing hormone progesterone, into an all-or-none biological response: oocyte maturation. The all-or-none character of the response can be accounted for in part by the intrinsic ultrasensitivity of the oocyte’s mitogen-activated protein kinase (MAPK) cascade. In contrast to the zero-order ultrasensitivity we implemented in the CM model above, here the steepness of the input-output curve originates mainly in the distributive nature of the phosphorylation events in the cascade [12].

We used a detailed, mass-action kinetics model of the MAPK cascade previously developed in a combined modeling-experimental study [13] (Figure 5A). This model describes the dynamics of 22 species participating in 10 reactions. The authors estimated or extracted from cellular and biochemical experiments each of the 37 parameters (See Supplementary Information) [13, 14]. We analyzed the shifter properties of this model. We simulated various input (E1_tot_) concentrations as steps and monitored the system output (doubly phosphorylated MAPK (MAPK-PP)) over time (Figure 5B). The output dynamics was sigmoidal, especially evident with low levels of input. That is, there was a long delay until output increased: MAPK-PP began to increase only after 20 minutes with E1 = 10^-5^. This suggests that transmission of information in this cascade is slow. Remarkably, its input-output curve shifted over time, and did so slowly, only reaching steady-state after about 100 minutes (Figure 5C). As with the case of our LR-CM model above, output was submaximal at short times even in the high input region. Interestingly, the ultrasensitivity of the system was not present at early times, but developed over time. This is similar to the behavior displayed by our CM cycle simulated in the zero-order regime (Figure 3I and L).

**Figure 5:**
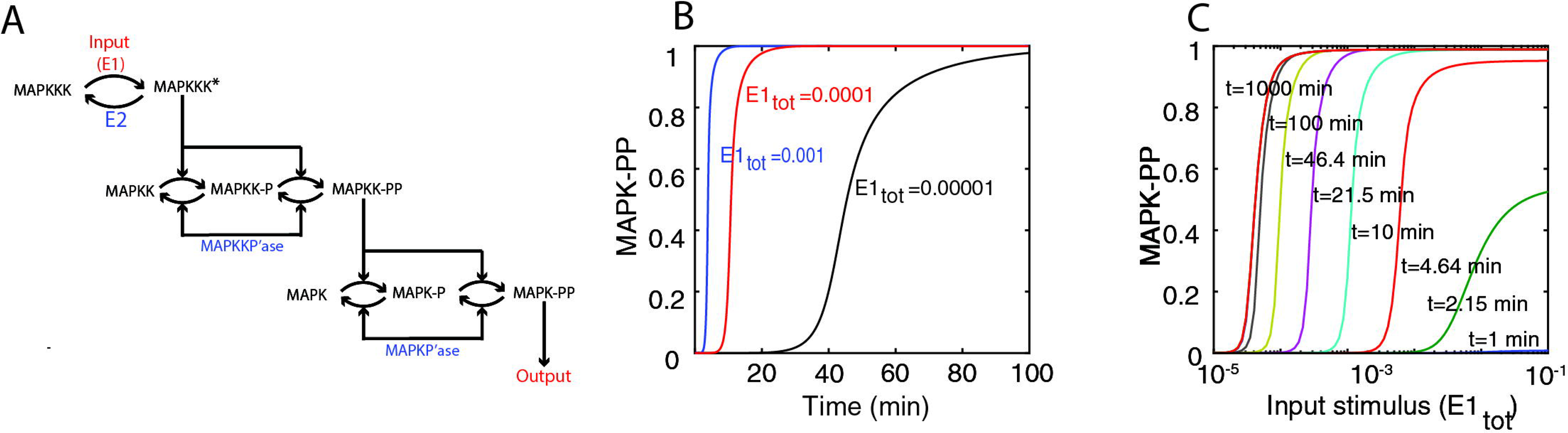
Pre-steady-state behavior in the oocyte maturation MAPK cascade model. **A**. Schematic view of the MAPK cascade. The cascade consists of a MAPK kinase kinase (MAPKKK), a MAPK kinase (MAPKK), and a MAPK. In this model, the dual phosphorylations of MAPK and MAPKK occur by a two-step, distributive mechanism. **B-C**. Simulation of the Huang & Ferrell model. Time evolution of normalized MAPK-PP for three different inputs and Input-output curves (normalized MAPK-PP versus E_1,tot_), for different times, as indicated.

In summary, our results indicate that cascades of covalent modification cycles are shifters. Therefore, the range of inputs that these systems would be able to distinguish will strongly depend on the timing of the input and of downstream outputs.

### Transcriptional regulation I. A simplified model of gene expression is not a shifter

Above we have studied how the shifting property propagates through a chain of simple biochemical reactions. However, signaling pathways usually lead to changes in gene expression. Thus, it is also interesting to consider whether a gene expression step could also be a shifter. As a first step, we considered the simplest case of a transcriptional regulatory network: a single transcription factor X induces the expression of gene *y* into protein Y at a rate γ by binding to a promoter region P in the DNA. Over time, Y is degraded at a rate d. This process may be described by the following scheme:

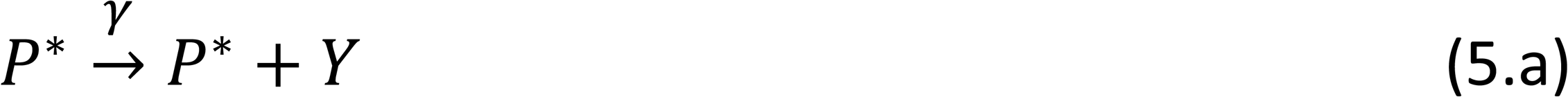

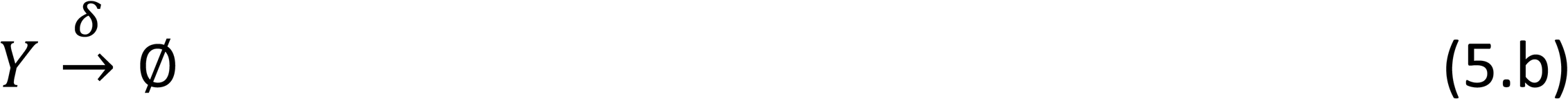

where P* stands for the active promoter. This translates into the following differential equation:

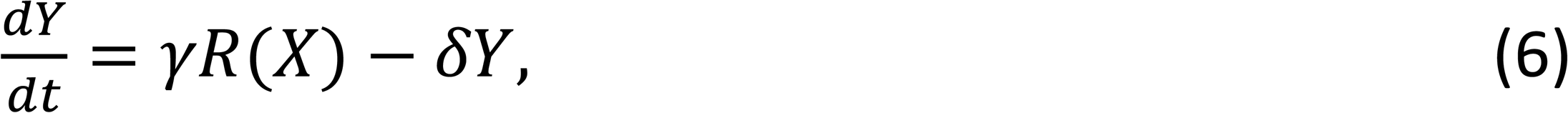

where R(X) is the regulatory function. Typically R(X) may take the form of a Hill function X^n^/(X^n^+θ^n^) with cooperativity n and EC_50_ indicated by θ, depending on the exact way in which X binds the DNA. The parameters γ and d are the maximum synthesis rate of protein Y and the degradation plus dilution parameter, respectively.

In the limit case in which X is not a function of time, equation (6) has a simple solution:

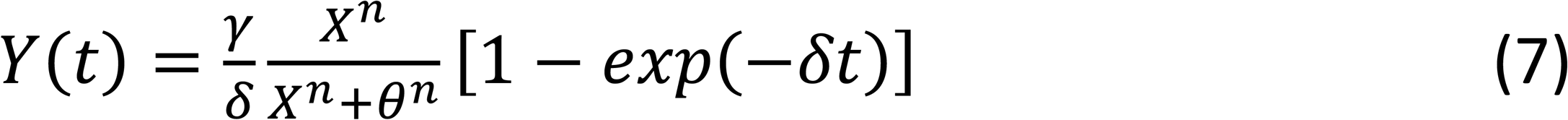

As has been previously noted, the dynamics of this system only depends on the degradation rate d, independently of the value of the stimulus X [15]. X only controls the amplitude of the response Y. Not only that, normalizing the input-output curve to the maximum at a given time, results in a time independent input-output curve:

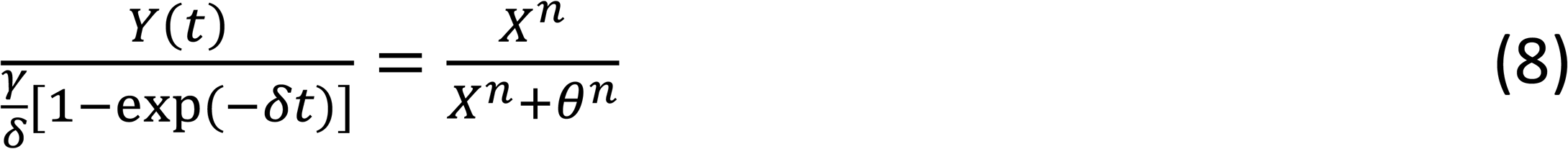

Therefore, this system is not a *shifter*, and as such may not operate in PRESS mode.

### Transcriptional regulation II. A more detailed model of gene expression is a shifter

We suspected that the inability of the above transcriptional model to behave as a shifter is not because transcription is intrinsically different from the other biochemical processes, but because the model is oversimplified, omitting one or more reactions crucial for this behavior. An example is the omission of binding between the transcription factor X and the promoter, since, as we explained above, the binding reaction is a shifter. Thus, we expected that a more detailed model of transcriptional regulation would behave as a *shifter*, at least in some region of the parameter space. To determine if that hypothesis was correct, we considered a model in which a transcription factor X binds a promoter P. The bound promoter Pb then transitions to an active promoter P*, which can then synthesize protein Y (Figure 6 Ai and Aii).

**Figure 6:**
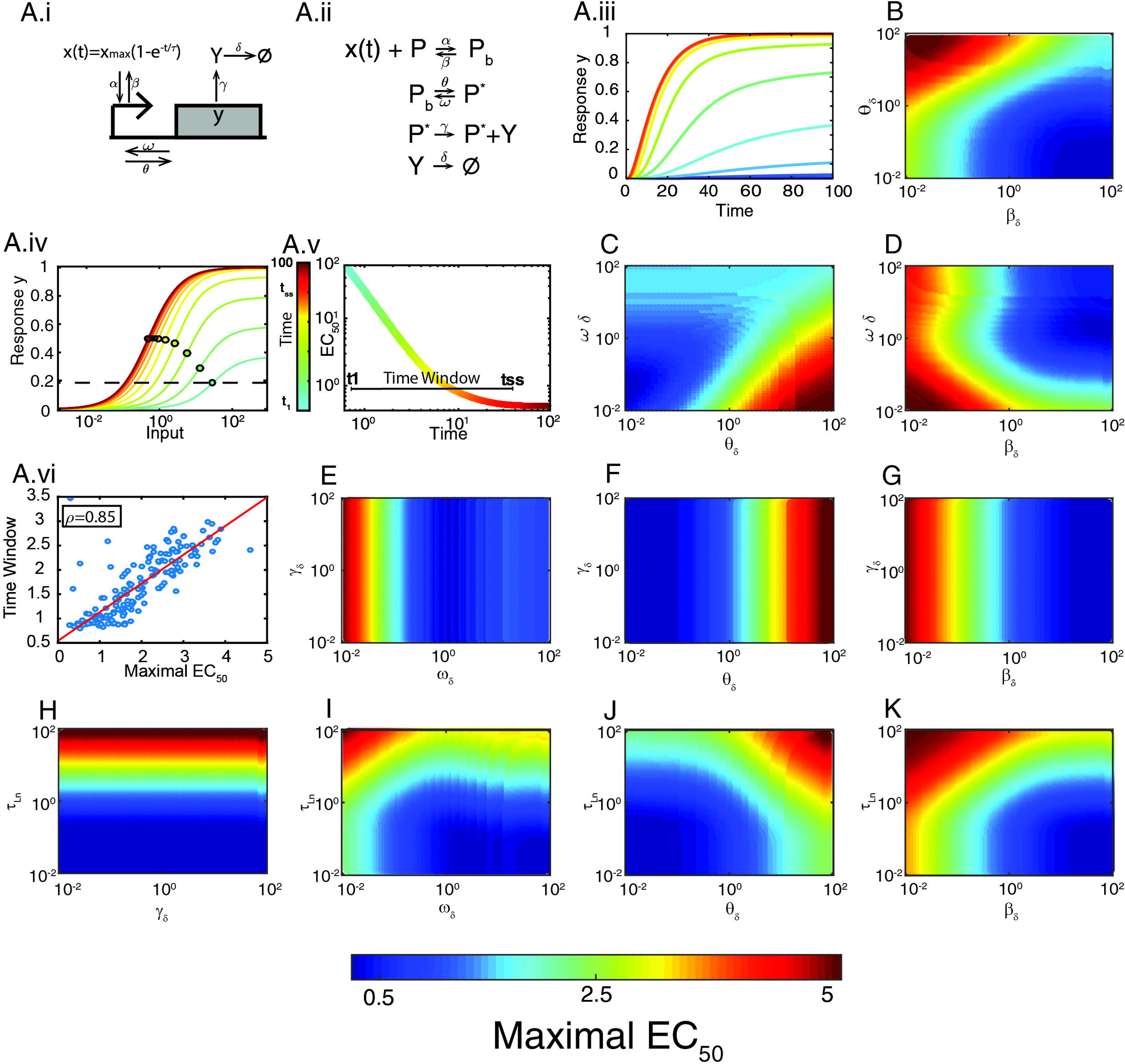
Shifting in a gene expression model. Diagram for transcription factor-mediated gene expression. X is the transcription factor; a filled rectangle represents gene y; the arrow pointing at the rectangle is the promoter region; α and β are the binding and unbinding rates of X to the promoter, θ and ω are the activation and deactivation rates of the promoter, γ and δ are the production and degradation+dilution rates for protein Y. **A.ii.** Reaction scheme of the model. **A.iii-A.iv.** Output Y vs time for different inputs (using a heatmap scale) and Output Y vs input X (concentration of the transcription factor) for different times (using a heatmap scale). Y has been normalized to its maximum (γ P_T_/d). Filled circles over the curves mark the EC_50_. Here all kinetic rates are equal to 1 and the activation time of X has a τ_n_ = 10. **A.v.** EC_50_ vs. time for the curves in **A.iv**. Maximal EC_50_ (ME) is indicated with a horizontal dotted line in **A.IV**, the Time window (TW) with a horizontal line in **A.v**. The color scale indicates the time. **A.vi.** Time window (TW) vs. Maximal EC_50_ (ME) for 200 randomly selected parameter sets out of the 2 10^5^ total scanned (ρ = 0.8562, using all parameter sets) (see SI for a description of the scanning procedure). **B-K**. Each plot shows ME (using a heatmap scale) for 150×150 values of the indicated parameters (see TW in Figure S2). In each panel, the rest of the parameters were equal to 1.

We described this system by the following equations:

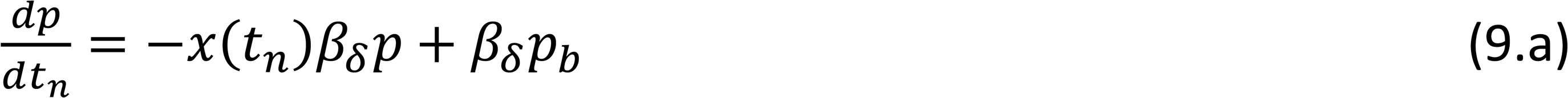

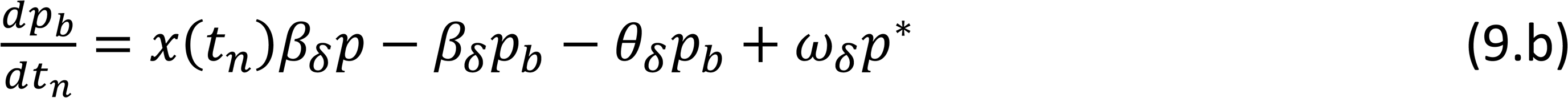

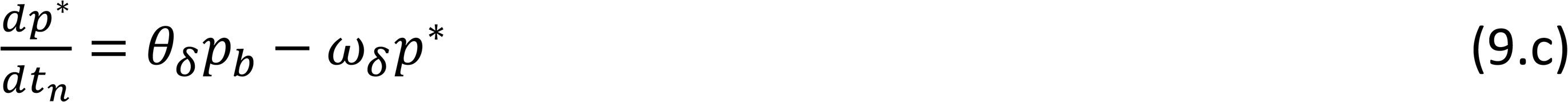

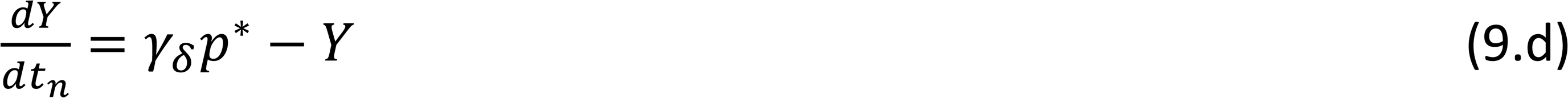

and the conservation law *p* + *p*_*b*_ + *p*^*^ *=* 1, where p, p_b_ and p* are the unbound, bound and active promoters, respectively, considering a system with one transcriptional unit (one promoter with one binding site), and the time is expressed in units of a reference time t_ref_=1/d, so that t_n_=t/t_ref_. This reference time is the time-scale of the degradation plus dilution processes. All the rates are in units of the degradation rate d, resulting in the following parameters *β*_*δ*_ = *β/δ, θ*_*δ*_ = *θ/δ, ω*_*δ*_ = *ω/δ*, *γ*_*δ*_ = *γT*_*T*_*/δ*.

Considering that the concentration of free X is not influenced by the binding and unbinding of transcription factor molecules (since there is only one site to bind), so that X ∼ X_T_. We described the dynamics of transcription factor accumulation by

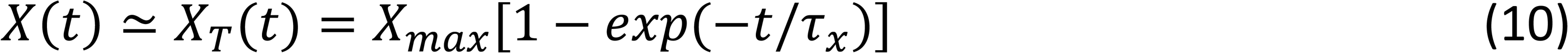

where X_max_ is the maximum value reached by X and τ_x_ is its characteristic time. In this way, and expressing the time in terms of the reference time *t*_*ref*_ *=* 1*/δ*, we obtained

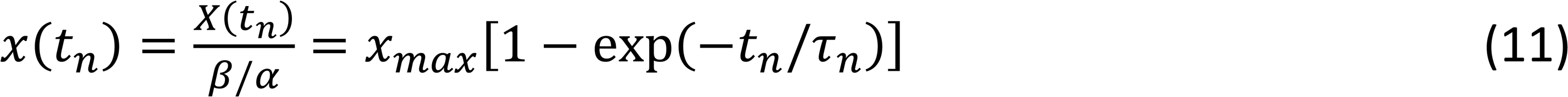

where *t*_*n*_ = *t/t*_*ref*_ and *τ*_*n*_ = *τ*_*x*_*/t*_*ref*_ are the dimensionless time and dimensionless accumulation time of X, respectively.

To analyze this model in terms of its kinetic parameters, we first studied its behavior (time course and input-output curve) using an arbitrary parameter set (Figure 6 A.iii and A.iv). Notably, the input-output curve shifted from right to left over time, indicating that this model of transcription does behave as a shifter. As in the cases of the LR-CM model (Figure 5), the amplitude of the curves changed as well. Thus, to analyze this type of models further, we defined a threshold output (20% of the maximal output possible) and the time at which it was obtained, t_1_. This time t_1_ defined the first moment in which there was a reasonable dose-response. We called the ratio EC_50_(t_1_)/EC_50_(t_ss_) of that curve Maximal EC_50_ (ME) (expressed in units of the EC_50_ at steady state). It was also useful to define the Time Window (TW) as the interval between t_1_ and the time the model reached steady-state (t_ss_):

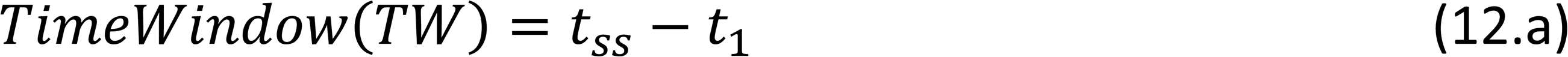

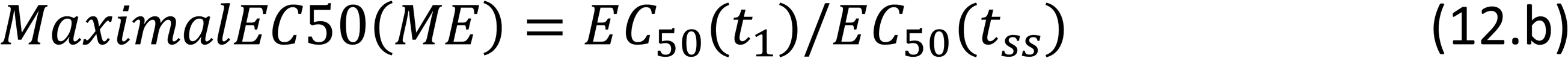

A high value of ME indicates that before steady state there is a wide range of distinguishable inputs, and a large TW indicates that there is a considerable time during which pre-steady state information could be influencing the system. Next, to determine how ME and TW depended on the parameters of the model, we varied two parameters at a time from 10^−2^ to 10^2^, while maintaining all the remaining parameters equal to the degradation rate *δ* (see Methods for a detailed description of the parameter scan and computation of TW and ME). In this way we generated 10^4^ sets for which we computed ME and TW (Figure 6 A.v). Several observations are worth mentioning. First, we noted that ME and TW were strongly correlated (Figure 6 A.iv), indicating that the slower the shifter, the larger the distance from the first observable EC_50_ and its steady state value. Second, ME/TW were independent of γ, the production rate of the protein Y (Figure 6 F-H). This was expected, since γ does not influence the dynamics of promoter activation and only appeared as a proportionality constant in the solution of the ODE for Y(t) (see equations 9 a-d). Third, ME/TW increased as either of the two rates that control inactivation of transcription (the dissociation rate of the transcription factor X (β) or the promoter deactivation rate (ω)) diminished (see for example Figure 6 D). Fourth, the activation (*θ*) and deactivation (ω) of the promoter were interdependent in the way they controlled ME/TW. When the ratio K = *ω/θ* was small, ME/TW was large (Figure 6 C). Finally, the rate of accumulation of active transcription factor X (*τ*_*n*_) affected the system as a whole, independently of all the other parameters: if *τ*_*n*_ ≫ 1, ME/TW was large, while if *τ*_*n*_ ≤ 1, then ME/TW was small.

In summary, the above results show that the process of gene expression behaves as a shifter. The shift was larger and it lasted longer if a) the active transcription factor increased slowly, and b) transcription inactivation was slow, both relative to the degradation rate of the gene product. Thus, systems with unstable gene products tend to be better shifters.

## Discussion

Previously we have shown that if the input-output curve of a ligand:receptor reaction is measured before it reaches equilibrium, it is *shifted* to the right of its equilibrium position (Fig 1C). More specifically, the EC_50_ decreases over time [4] (Fig 1E). In this work we refer to systems with this property as *shifters*. Shifting is based on a particular feature of the system: it reaches equilibrium with a characteristic time that is a decreasing function of the input (Fig 1D). That is, ligands need appreciably more time to reach equilibrium binding at low concentrations than at higher concentrations. Thus, shortly after stimulation, binding is further from equilibrium binding the lower the ligand concentrations. Consequently, at that early time, while the amount of response may already be close-to-steady-state at the highest ligand concentrations, it may still be far less than expected at steady-state at the lowest concentrations. This will result in a steeper saturation curve (i.e. nH > 1; Fig 2E) and the half-maximal binding (EC_50_) (Fig 2E) values may also be appreciably higher than expected at a steady-state. We have also shown that *shifting*, coupled to a downstream pathway appropriately fast and transient, allows systems to use pre-equilibrium information in order to distinguish stimuli that saturate receptors at equilibrium. We called this ability PRESS (Pre-Equilibrium Sensing and Signaling) [4].

In our previous article [4], the speed of the *shift* depended exclusively on the binding/unbinding rates: the slower the rates, the slower the *shift* and the longer the time window during which the system may exploit pre-equilibrium information. Thus, we speculated that biological systems exposed to step-like stimulus might have evolved slow binding ligand-receptor pairs to extend the range of inputs it can distinguish (the dynamic range). However, in many instances, stimuli increase slowly. Thus, in this work we asked if the requirement for PRESS that the binding reaction be slow may be relaxed by introducing dynamics to the stimulus. Indeed, here we show that input dynamics controls the speed of the *shift*.

The other important goal of the current paper was to find other basic signaling systems that could work as *shifters*, apart from a ligand-receptor binding. We found that covalent modification cycles and simple gene expression systems can *shift* their input-output curve in time. In each case, we studied which conditions improve the *shifting* capacities. The motivation for testing covalent modification cycles as *shifters* comes from the fact that when the enzymes operate in the first order regime, these cycles follow the same mathematical description that is valid for a ligand-receptor system [16]. Thus, at least in that limit, a decreasing EC_50_ versus time was guaranteed. Simple gene expression systems, on the contrary, were not expected to work as *shifters* because in this case the response time is governed by the degradation/dilution rate only [17], which is usually independent on the input (e.g. transcription factor level) received by this system. However, we found that only an oversimplified model of gene expression has no *shift*. As soon as the model incorporated additional, more realistic reactions, such as transcription factor binding to the promoter, shifting emerged.

### Ligand-receptor systems

When receptor binding/unbinding is slow, the EC_50_ changes correspondingly slowly. Here, we asked how the *shift* in input-output curves is altered when the input has dynamics, particularly when ligand accumulation is slower than the binding/unbinding reaction speed. We considered the case of a ligand that accumulates exponentially. We focused on the impact that the relative time-scales (ligand accumulation versus binding/unbinding) could have on the *shift*. We found that a slow rising input caused a slow *shift*, providing a longer time to a pathway downstream to operate in PRESS mode. Thus, if the stimulus is slow in rising, then the *shift* will also be slow, even in cases where the ligand-receptor interaction is fast. For the case in which ligand accumulation is much slower than the binding/unbinding reaction, we derived a quasi-steady-state approximation leading to an analytical expression for the temporal profile of the EC_50_. Interestingly, in this limit the slowness of the *shift* is directly controlled by the slowness of ligand accumulation.

### Covalent modification cycle

In this work we studied CM cycles in their ability to behave as *shifters*, assuming that the activating enzyme increases following an exponential function, for example due to its accumulation after its expression is induced. We considered two extreme regimes: the first-order and the zero-order regimes, both for the cases in which the activating enzyme accumulates fast or slow relative to the timescale of the covalent modification cycle. Mixed scenarios, like saturated kinase and non-saturated phosphatase, and vice versa, are for sure of interest and could lead to different operating regimes [18] but were not covered in the present article.

As obtained for the RL system, we found that when the stimulus increased slowly, the leftward *shift* in the input-output curve was correspondingly slow. We also found that the *shift* was faster in zero-order, indicating that there is less time for PRESS when the enzymes are saturated.

The first-order regime of a CM cycle, mathematically, is similar to the RL system. Thus, we expected that both the EC_50_ and n_H_ behaved over time similarly as well. Although the dynamics of the EC_50_ does indeed behave the same, the dynamics of the n_H_ was slightly different: it does not decrease monotonically form a 1.42 value towards 1 at steady-stete, but it increases from a value close to 1, peaks at 1.42 and only then it declines towards 1.

In the zero-order regime, we found that the ultrasensitivity evidenced in its high n_H_ takes time to develop, increasing from about 1 towards its final high steady-state value. This surprising result is due to the fact that the system is not actually in the zero-order for backwards reaction at early times, since at these times there is not yet product formed (the substrate for the backwards reaction). Thus, at early times, the system is in first order for one reaction and zero-order for the other.

### Coupling shifters

We considered the *shifting* properties of two coupled systems, a ligand-receptor coupled to a CM cycle, and a cascade of CM cycles. For the former, we evaluated the *shifting* property throughout the parameter space; for the later, we analyzed a model developed to capture the concrete case of a MAPK cascade involved in oocyte maturation. We found that the LR-CM system as a whole behaves as a *shifter* and also that the *shifting* of the coupled system is slower than that of each step alone. Regarding the MAPK cascade, we found a very slow *shift*. As found for a single CM cycle, the ultrasensitivity of a cascade of CM cycles develops over time and thus threshold input required to activate the switch moves over time. In summary, our results indicate that cascades of covalent modification cycles are *shifters*. Therefore, the range of inputs that these systems would be able to distinguish will strongly depend on the timing of the input and of downstream outputs.

### Transcriptional regulation

We found that this system is not a *shifter* only in the limit case in which its dynamics simply depends on the degradation rate. The reason for this is that this limit case omits reactions that are crucial for the shifting behavior, like the binding between the transcription factor and the promoter. A more detailed model for transcriptional regulation did behave as a *shifter*. The *shift* was larger and it lasted longer if a) the active transcription factor increased slowly, and b) transcription inactivation was slow, all this relative to the degradation rate of the gene product. Thus, systems with unstable gene products tend to be better *shifters*.

### Shifting mechanism might be widespread

Summarizing, under physiological conditions, receptor binding and activation, and subsequent signaling are often dependent on very short stimuli and usually do not reach a steady-state. Therefore, it is relevant to study receptor activation and signaling, its potencies, efficacies and sensitivities under non-equilibrium conditions.

Our results here apply particularly to systems that need to adapt their input dynamic range (the range of inputs for which the system is able to generate distinguishable outputs) according to different scenarios [4]. Binding/unbinding reactions, covalent modification cycles and gene expression regulation are the building blocks of cellular regulation, underlying all processes in the cell, from basic homeostatic maintenance of cellular structures and energy balance, to cell division, cell differentiation. The fact that we have shown that these elements all may behave as shifters indicates that attention to dynamics of each of these processes might shed interesting insights into fundamental biological processes.

## Methods

### Numerical integration of ODEs

ODEs were integrated using the ode23s function from Matlab.

### Covalent-Modification cycle: parameter scanning

We scanned parameter values randomly with log uniform distribution within the intervals defined on Table S1. We used only random parameter sets whose *K*_*a*_ and *K*_*d*_ fell within the intervals.

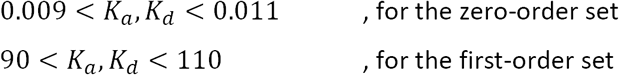

*K*_*a*_ and *K*_*d*_ correspond to the Michaelis-Menten constants *K*_*a*_ *=* (*d*_1_ + *k*_1_)*/a*_1_*S*_*T*_, *K*_*d*_ *=* (*d*_2_ + *k*_2_)*/a*_2_*S*_*T*_ in units of the total amount of substrate. We kept the first 100 sets satisfying the mentioned conditions. We then simulated the CM cycle using each of those sets and obtained input-output curves for 50 different times within the interval [10^−3^,10^3^], for fast and slow input.

### A ligand-receptor activates a covalent modification cycle: parameter scanning

To obtain different parameter sample sets, we used the same criterion as with the CM cycles. Time was in units of *k*_*off*_ (i.e. parameters are in units of the unbinding rate) and the input in units of the dissociation constant of the binding reaction. We scanned *a*_1_, *a*_2_, *d*_1_, *d*_2_, *k*_1_, *k*_2_, *S*_*T*_ and *E*_*T*_ randomly, from a uniform distribution within the intervals defined in Table S1, and R_T_ in the interval [01500]. This interval was chosen based on the total amount of substrate (see Table S1) to cover three possible scenarios: *R*_*t*_ ≪ *S*_*T*_, *R*_*t*_ ∼ *S*_*T*_, *R*_*t*_ ≫ *S*_*T*_). Under these considerations, we kept two groups of parameter sets that satisfied

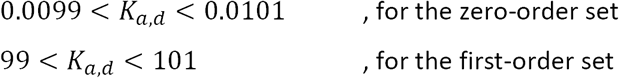

In Figure 4 we show the behavior of 200 parameter sets for each group that produced an output *S*^*^ with an amplitude greater than 0.1 (10% of S_T_).

### Transcriptional regulation

#### 1) Parameter scanning

To quantify the effect that each parameter has on the shifter, we scanned parameters *β*_*δ*_, *θ*_*δ*_,*ω*_*δ*_, *γ*_*δ*_, and *σ*_*δ*_ with the following approach. We varied two parameters in a grid of 150 × 150 in log scale from 10^−2^ to 10^2^ while maintaining the remaining parameters fixed to unity (i.e., kinetic rates and inverse time scale *σ* were equal to the degradation rate *δ*). For each point on the grid, we computed the Maximal EC_50_ and the Time Window, as defined in the main text. We studied all possible pair of parameters.

#### 2) Computing *t*_1_ and *t*_*ss*_

To compute the first observable time, we defined t_1_ as the time that satisfies

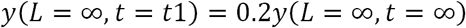

the time at which the (dimensionless) output reached 20% of the maximal output. We obtained this value *t*_1_ using a time grid of *N*_*T*_ *=* 1000 and varying the time in linear scale from 0 to 50 (time in dimensionless units).

Having t_1_, we computed dose-response curves for different times. We varied the dose in log scale from 10^−2^ to 10^4^ with 50 values, and time in log scale from *t*_1_ to 10^3^ using 500 values. Such a density in the time grid is needed to accurately compute EC50 versus time.

We obtained *t*_*ss*_ as the first time at which 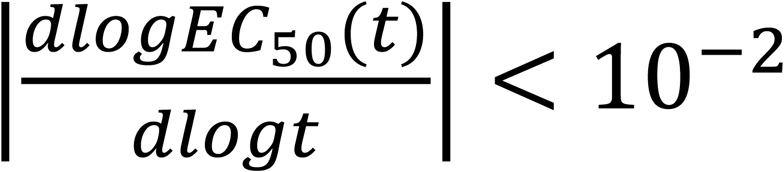. As *EC* (*t*) decreases monotonically with an asymptotic value, its log derivative is negative and goes asymptotically to zero. Hence, we considered that the dose response curve reached steady state when 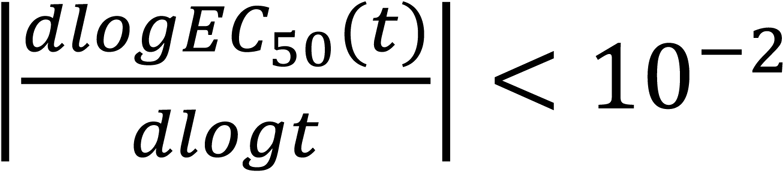. Note that, strictly, steady state is reached at different times for different inputs. However, for practical reasons, we used the time that the EC_50_ reached steady state as a proxy for the whole input-output curve. Having *EC*_50_(*t*), *t*_1_, and *t*_*ss*_, we used equation 12 to compute Maximal EC_50_ and Time Window.

## Acknowledgements

This work was supported by a grant from the from the Argentine Agency of Research and Technology (PICT2013-2210) to ACL and ACV.

## Author contributions

ACV and ACL designed the project. JDB and ACV performed all mathematical analysis and simulations. JDB, ACV and ACL analyzed the results and prepared the manuscript.

